# A single nucleotide variant in the PPARγ-homolog *Eip75B* affects fecundity in *Drosophila*

**DOI:** 10.1101/2021.12.07.471536

**Authors:** Katja M Hoedjes, Hristina Kostic, Thomas Flatt, Laurent Keller

## Abstract

Single nucleotide polymorphisms are the most common type of genetic variation, but how these variants contribute to the evolutionary adaptation of complex phenotypes is largely unknown. Experimental evolution and genome-wide association studies have demonstrated that variation in the PPARg-homolog *Eip75B* is associated with longevity and life-history differences in the fruit fly *Drosophila melanogaster*. Using RNAi knockdown, we first demonstrate that reduced expression of *Eip75B* in adults affects lifespan, egg-laying rate and egg volume. We then tested the effect of a naturally occurring SNP variant within a cis-regulatory domain of *Eip75B* by applying two complementary approaches: a Mendelian randomization approach using lines of the *Drosophila* Genetic Reference Panel, and allelic replacement using precise CRISPR/Cas9-induced genome editing. Our experiments reveal that this natural polymorphism has a significant pleiotropic effect on fecundity and egg-to-adult viability, but not on longevity or other life-history traits. These results provide a rare functional validation at the nucleotide level and identify a natural allelic variant affecting fitness and life-history adaptation.

## INTRODUCTION

Single nucleotide polymorphisms (SNPs) are the most abundant form of genetic variation and are an important source of fuel for evolutionary adaptation (Suh and Vijg 2005). Advances in high-throughput genome sequencing technologies have amounted to the association of vast numbers of natural variants with complex, quantitative traits (Schlötterer, et al. 2014), but identifying the true causal variants remains a challenge as loci within a genomic region are often genetically linked to each other, resulting in false positive signals (Franssen, et al. 2015; Barghi and Schlötterer 2019). Therefore, functional characterization of individual candidate loci is critical to pinpoint the mechanisms of adaptation and quantify the phenotypic impact of these loci.

Homologous allele replacement at the endogenous locus is the ‘gold standard’ for demonstrating a causal effect at base-pair resolution, but such functional tests are rare, especially for multicellular organisms (Turner 2014; Zdraljevic, et al. 2017; Ramaekers, et al. 2019; Zdraljevic, et al. 2019; Na, et al. 2020; Vigne, et al. 2021). Instead, functional analyses are often performed at the gene level by generating (null) mutants or using RNAi (Zhou, et al. 2011; Martins, et al. 2014; Fabian, et al. 2018; Huang, et al. 2020; Parker, et al. 2020). Functional effects of natural alleles may, however, be more subtle or different altogether from mutant alleles (Stern 2011; Mokashi, et al. 2021; Hoedjes, et al. 2022). The aim of this study was to conduct functional analyses of a natural polymorphism of *Ecdysone-induced protein 75B (Eip75B*), a candidate gene for ageing and life-history evolution in fruit flies.

*Eip75B* encodes a nuclear hormone receptor homologous to peroxisome proliferator-activated receptor gamma (PPARg), a key regulator of adipogenesis and pregnancy-induced modifications in lipid metabolism in mice and humans (Waite, et al. 2000; Hong and Park 2010; Zipper, et al. 2020). Nuclear hormone receptors, such as *Eip75B*,are evolutionarily conserved regulatory hub genes that integrate environmental factors, hormonal signaling, and metabolism and relay these inputs to downstream targets (Palanker, et al. 2006; Jagannathan and Robinson-Rechavi 2011; Hoffmann and Partridge 2015). In the fruit fly, *Eip75B* also acts as an early-response gene in the ecdysone signaling pathway, which plays a critical role in the regulation of development, physiology and lifehistory of arthropods, similar to thyroid hormones in vertebrates (Bialecki, et al. 2002; Simon, et al. 2003; Flatt, et al. 2006; Terashima and Bownes 2006; Toivonen and Partridge 2009; Magwire, et al. 2010; Caceres, et al. 2011; Galikova, et al. 2011; Yamanaka, et al. 2013; Belles and Piulachs 2015; Jaumouillé, et al. 2015).

*Eip75B* has been identified as an “ageing” candidate gene by two independent genome-wide association studies (Huang, et al. 2020; Pallares, et al. 2020) and three experimental evolution experiments (Carnes, et al. 2015; Hoedjes, et al. 2019; Parker, et al. 2020), but each study detected different candidate SNPs. Experimental evolution of longevity has been intricately linked to changes in fecundity, suggesting some shared genetic basis of these two traits possibly due to antagonistic pleiotropy (Remolina, et al. 2012; Carnes, et al. 2015; Fabian, et al. 2018; May, et al. 2019; Flatt 2020). Moreover, evolutionary changes in longevity can be associated with changes in viability, development time, immune function, stress resistance, and even learning ability (e.g., Chippindale, et al. 1994; Mery and Kawecki 2005; Baldal, et al. 2006; Fabian, et al. 2018; Flatt 2020). Given its central role in hormonal signaling and its pleiotropic effects on numerous developmental and physiological processes, *Eip75B* may thus be involved in the concerted evolution of longevity and correlated life-history traits.

In this study, we first used RNAi silencing of *Eip75B* in adults to determine the effects of the gene on lifespan, egg laying and egg volume. We then selected a naturally occurring SNP in a cis-regulatory domain of *Eip75B* that was previously identified as a candidate in an experimental evolution study (Hoedjes, et al. 2019), and functionally characterized it using a SNP-specific association study based on Mendelian randomization and precise allelic replacement using CRISPR/Cas9. Our results demonstrate that a single nucleotide variant can have a major effect on an important fitness-related trait, namely fecundity.

## RESULTS AND DISCUSSION

### RNAi knock-down of *Eip75B* in adults reduces longevity, egg-laying rate and egg volume

Recent studies have found that weak, constitutive knockdown of *Eip75B* throughout development and the adult stage increases lifespan in both males and females (Huang, et al. 2020), whereas tissue-specific knockdown in the amnioserosa, ovaries and accessory glands causes decreased lifespan but increased fecundity in females (Parker, et al. 2020). Given that *Eip75B* might have potentially confounding pleiotropic developmental effects on adult life-history, we investigated the effect of gene knockdown in adults only. The effects of *Eip75B* knockdown on longevity, fecundity (measured as the daily egg-laying rate per female) and egg volume were assessed with four independent RNAi constructs expressed in the adult stage using the mifepristone-inducible, constitutive *daughterless-GS-GAL4* driver (Osterwalder, et al. 2001). RT-qPCR confirmed a significant, partial knockdown of *Eip75B* for three of the four constructs, E01, E03, and E44, but not for the fourth construct (E10) (Supplementary Table 1). The fourth construct was nonetheless included in the study as it had been observed to affect lifespan and fecundity in previous studies (Huang, et al. 2020; Parker, et al. 2020).

Knockdown of *Eip75B* expression resulted in a significant reduction in lifespan only for constructs E01 (females, up to −15.9%) and E44 (both sexes; females: up to −24.5%, males: up to −18.2%) as compared to the control, although we also observed a similar trend in lifespan reduction for construct E03 (Figure 1A). The effects of *Eip75B* knockdown on egg-laying rate and egg volume were more consistent, with a significant reduction in both egg-laying rate and egg volume for constructs E01, E03, and E44 (a reduction of the total number of eggs laid per female of 27.1%, 23.2%, and 57.7%, respectively; and a reduction in egg volume of up to 6.6%, 28.8%, and 23.4%, respectively), and a weak trend for construct E10 (Figure 1B and 1C).

**Figure 1:**
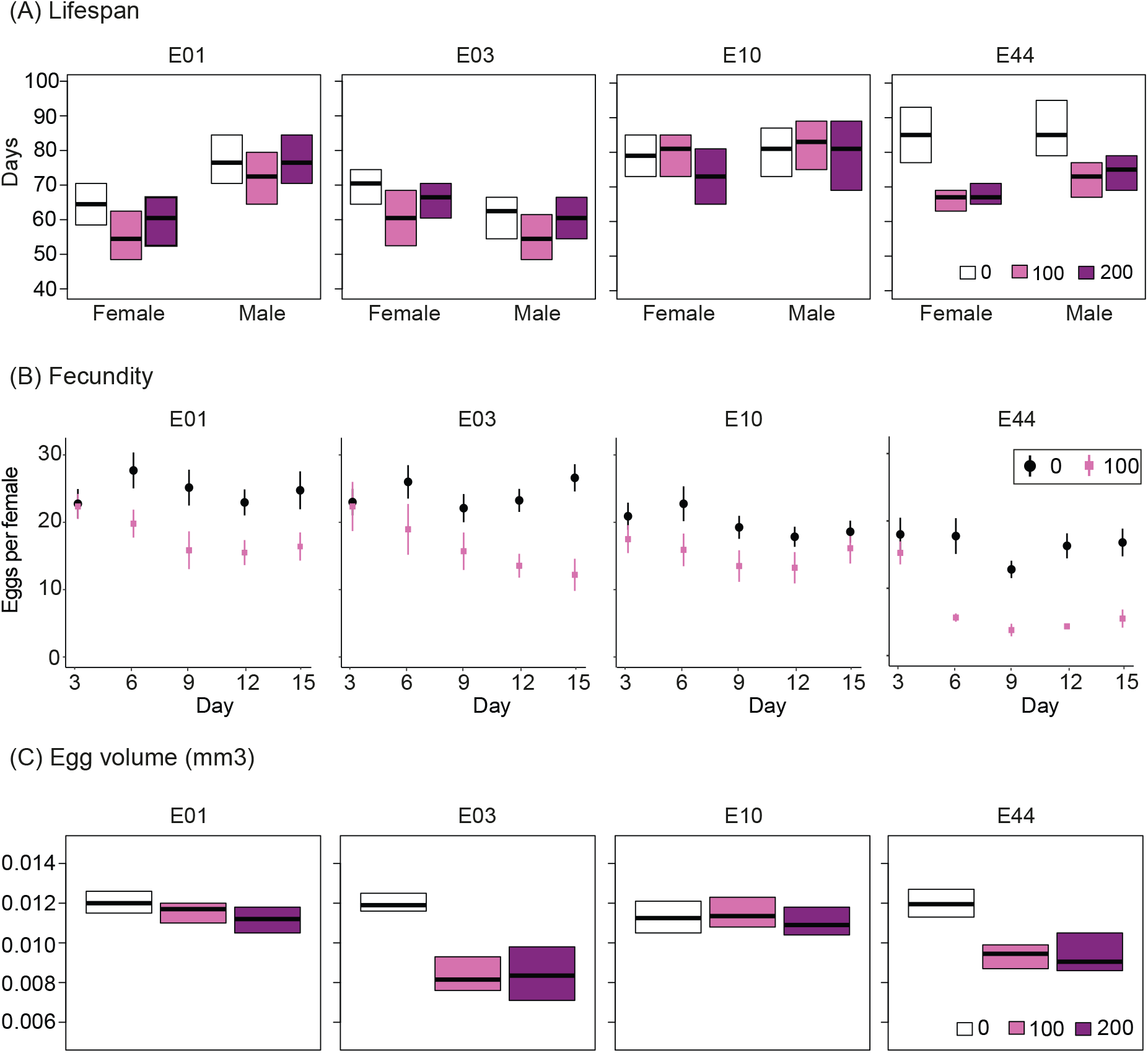
Effects of RNAi knockdown of *Eip75B* on lifespan, fecundity, and egg volume. Effects of RNAi on *Eip75B* in the adult stage were assessed using four different UAS-RNAi constructs (E01, E03, E10, and E44) under control of the mifepristone-inducible *da*-GS-GAL driver. The effects of mifepristone concentration (0, 100, or 200 μg/ml), as measure for RNAi knockdown, on lifespan were considered significant when P<0.0042 (Bonferroni: 0.05/12), whereas effects on egg-laying rate and egg volume were considered significant when P<0.01 (Bonferroni: 0.05/5); asterisks indicate significant results. (A) *Eip75B* knockdown resulted in a significant reduction in lifespan for the constructs E44 (female: χ^2^ = 22.7, *P* < 0.0001, male: χ^2^ = 12.3, *P* = 0.0021) and E01 (female: χ^2^ = 16.9, *P* = 0.0002, male: χ^2^ = 3.0, *P* = 0.22), but not E03 (female: χ^2^ = 1.7, *P* = 0.44, male: χ^2^ = 4.5, *P* = 0.1) and E10 (female: χ^2^ = 5.3, *P* = 0.071, male: χ^2^ = 2.6, *P* = 0.28). (B) *Eip75B* knockdown had a significant effect on fecundity for constructs E01 (χ^2^ = 20.4, *P* < 0.0001), E03 (χ^2^ = 64.7, *P* < 0.0001) and E44 (χ^2^ = 96.8, *P* < 0.0001), but not E10 (χ^2^ = 3.5, *P* = 0.18). (C) Also, *Eip75B* knockdown had a significant effect on egg volume for constructs E01 (*F* = 6.3, *P* = 0.0028), E03 (*F* = 68.7, *P* < 0.0001) and E44 (*F* = 54.1, *P* < 0.0001), but not E10 (*F* = 2.8, *P* = 0.066).

Our results provide a functional validation at the gene level, confirming a potential role of *Eip75B* in affecting lifespan and fecundity, in line with previous studies (Huang, et al. 2020; Parker, et al. 2020). However, comparing our data with these previous studies also suggests that the direction and the magnitude of the observed phenotypic effects can depend on the timing, the tissue, and the type of genetic modification. This indicates that functional tests at the level of an entire gene are useful to reveal potential functions of a gene, but that they may not provide an accurate estimation of the exact functional impact of its allelic variants. Functional tests at the nucleotide level are, therefore, necessary to understand to role of *Eip75B* in the evolution of ageing and life history.

### Natural polymorphisms at *Eip75B* have been linked to variation in longevity and fecundity

The *Eip75B* SNPs which have been identified as potentially affecting ageing and life history are located throughout the gene, but mostly in non-coding regions, and may, therefore, have regulatory effects (Turner, et al. 2011; Carnes, et al. 2015; Jha, et al. 2015; Hoedjes, et al. 2019; Huang, et al. 2020; Pallares, et al. 2020; Parker, et al. 2020). *Eip75B* is a large (>100 kb) gene, comprising seven transcript variants that are expressed from alternative promotors. Isoforms have different temporal and spatial expression patterns and fulfill different, even opposing, functions in insects (Keshan, et al. 2006; Terashima and Bownes 2006; Bernardo, et al. 2009; Cruz, et al. 2012; Li, et al. 2016; Zipper, et al. 2020). Given this complex genomic architecture and the pleiotropic effects of *Eip75B*, independent SNPs may have different regulatory functions.

We considered a set of 16 SNPs which had been suggested to influence several life history traits, including longevity and fecundity, in an experimental evolution study in which flies where selected to reproduce at different ages (Hoedjes, et al. 2019; May, et al. 2019). From this set, we selected the most promising SNP (3L:18026199, genome release 6) for functional testing, based on its location within an inferred cis-regulatory motif (REDfly database v.9.5.1; Rivera, et al. 2019) and because it was among the three most significantly diverged SNPs. This SNP is bi-allelic, with the “T” variant being more common in the late-reproducing, long-lived populations (average frequency: 0.84 “T”, 0.16 “G”) as compared to the early-reproducing populations (average frequency: 0.52 “T”, 0.48 “G”) (Hoedjes, et al. 2019). A population genetic analysis revealed that these frequencies fall within the range of frequencies observed in natural populations of *D. melanogaster* in Europe and North America, suggesting that this SNP may also play a role in the adaptation of longevity and/or reproduction in natural populations (Supplementary File 1).

### A regulatory *Eip75B* SNP is associated with variation in fecundity and viability

We applied two independent genetic approaches to functionally test if our candidate *Eip75B* SNP affects lifespan and/or fecundity. First, we used a genetic association study based on the principle of “Mendelian randomization” (e.g., Glaser-Schmitt and Parsch 2018; Durmaz, et al. 2019; Betancourt, et al. 2021; Glaser-Schmitt, et al. 2021). This approach aims to identify potentially causal loci by testing the different genotypic states in a genetically diverse background, which limits potentially confounding (epistatic) effects associated with specific backgrounds. Here, we assessed a set of F1 crosses of various lines of the *Drosophila* Genetic Reference Panel (DGRP) (Mackay, et al. 2012) differing in their genotype (i.e. “TT” vs. “GG”). This heterozygous F1 approach was chosen as the DGRP strains are inbred and may thus suffer from reduced fitness due to recessive deleterious alleles. To test for the presence of potentially confounding factors of the genetic background, we analyzed linkage disequilibrium (LD, as measured by pairwise *r*^2^) among *Eip75B* SNPs of the DGRP panel, and estimated genetic differentiation (SNP-wise *F_ST_*) among the panels of F1 crosses with different genotypes. These analyses indicated that no other SNPs in the genome are in strong LD with our candidate SNP, which limits the chance that they might have a confounding effect on lifespan or fecundity (Figure 2A, Supplementary File 2).

**Figure 2:**
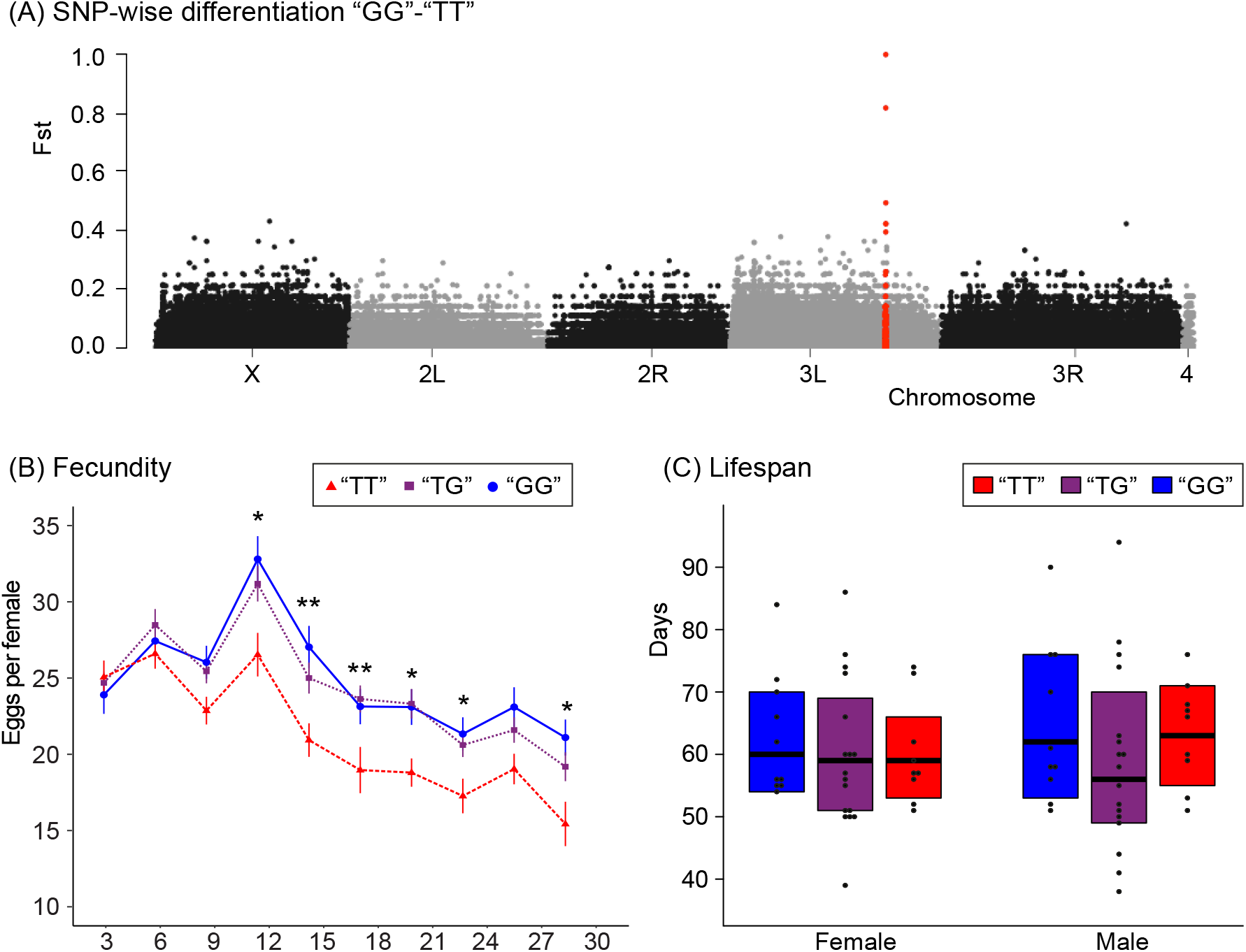
Mendelian randomization approach to test the association of an *Eip75B* candidate SNP with lifespan and fecundity. Correlation of the “T” versus “G” identity of the biallelic *Eip75B* SNP at position 3L:18026199 with fecundity and lifespan was tested by measuring F1 crosses of the *Drosophila* Genetic Reference Panel. (A) To test for potentially confounding factors in the genetic background, calculated SNP-wise genetic differentiation (*F_ST_*) using the pooled genome sequence information of all lines of the two homozygous sets of crosses (i.e. “TT” *versus* “GG”). These analyses indicate that only the candidate SNP is fixed (*F_ST_* = 1) between the two panels, and that very few SNPs show a high differentiation (*F_ST_* > 0.5) between the two panels, suggesting little or no confounding genetic factors. All *Eip75B* SNPs are indicated in red. (B) There was a significant effect of SNP genotype on fecundity (genotype: χ^2^ = 53.6, *P* <0.0001, genotype*day: χ^2^ = 45.3, *P* <0.0001). Asterisks indicate significant effects of genotype per day. Tukey post-hoc tests indicated that the “GG” and “TG” genotype had a similar fecundity on all days tested (*P*>0.05), whereas the “TT” and “TG” differed significantly in their fecundity on day 15 (*P*=0.028), day 18 (*P*=0.006, days 21 (*P*=0.008), and days 24 (*P* =0.041). (allele: χ^2^ = 50.9, *P*< 0.0001, allele*day: χ^2^ = 44.0, *P* <0.0001). (C) There was no significant effect of SNP genotype on lifespan (female: χ^2^ = 1.3, *P* = 0.51; male: χ^2^ = 1.7, *P* = 0.42).

F1 crosses homozygous for the “G” allele (and with a diverse genomic background) showed a significantly and consistently higher egg-laying rate than crosses homozygous for the “T” allele (Figure 2B), resulting in a 17.8% higher total number of eggs laid per female. F1 crosses that were heterozygous (“TG”) had an egg-laying rate that was not statistically different compared to crosses that were homozygous for the “G” allele, but higher (+14.8%) than crosses homozygous for the “T” allele for most of the time points assessed (Figure 2B). This suggests dominance or partial dominance of the “G” allele over the “T” allele for fecundity. On the other hand, there was no significant association between the candidate SNP and longevity. These findings suggest that the “G” allele might have played a role in facilitating higher (early) fecundity of the populations selected for early reproduction, but that this SNP is probably not causally responsible for the differences in lifespan and late reproduction observed in the experimental evolution experiment (May, et al. 2019). Given the lifespan effects of silencing *Eip75B*, other, as of yet unidentified alleles of *Eip75B* might play a causal role in longevity. Moreover, potentially confounding epistatic effects of the genetically diverse background could not be ruled out completely with our SNP association approach (Mokashi, et al. 2021).

Next, we used CRISPR/Cas9-induced genomic editing to modify the candidate SNP in a single genetic background without introducing any markers or off-target modifications that might interfere with gene function in the genomic region surrounding the focal SNP. We constructed two edited lines homozygous for the “G” allele and two lines homozygous for the “T” allele and tested their longevity, fecundity, and other life-history traits that *Eip75B* is known to affect. The lines carrying the “G” allele exhibited a significantly higher egg-laying rate (19.6% higher total number of eggs per female) and higher egg-to-adult viability (+33.3%) than lines with the “T” allele (Figure 3). Lines with the “G” allele also showed a small but significant reduction in starvation resistance in females (−8.5%). This indicates that this SNP has multiple pleiotropic functions. No significant differences between the two allelic states were observed for lifespan, egg-to-adult development time, adult dry weight, and egg volume.

**Figure 3:**
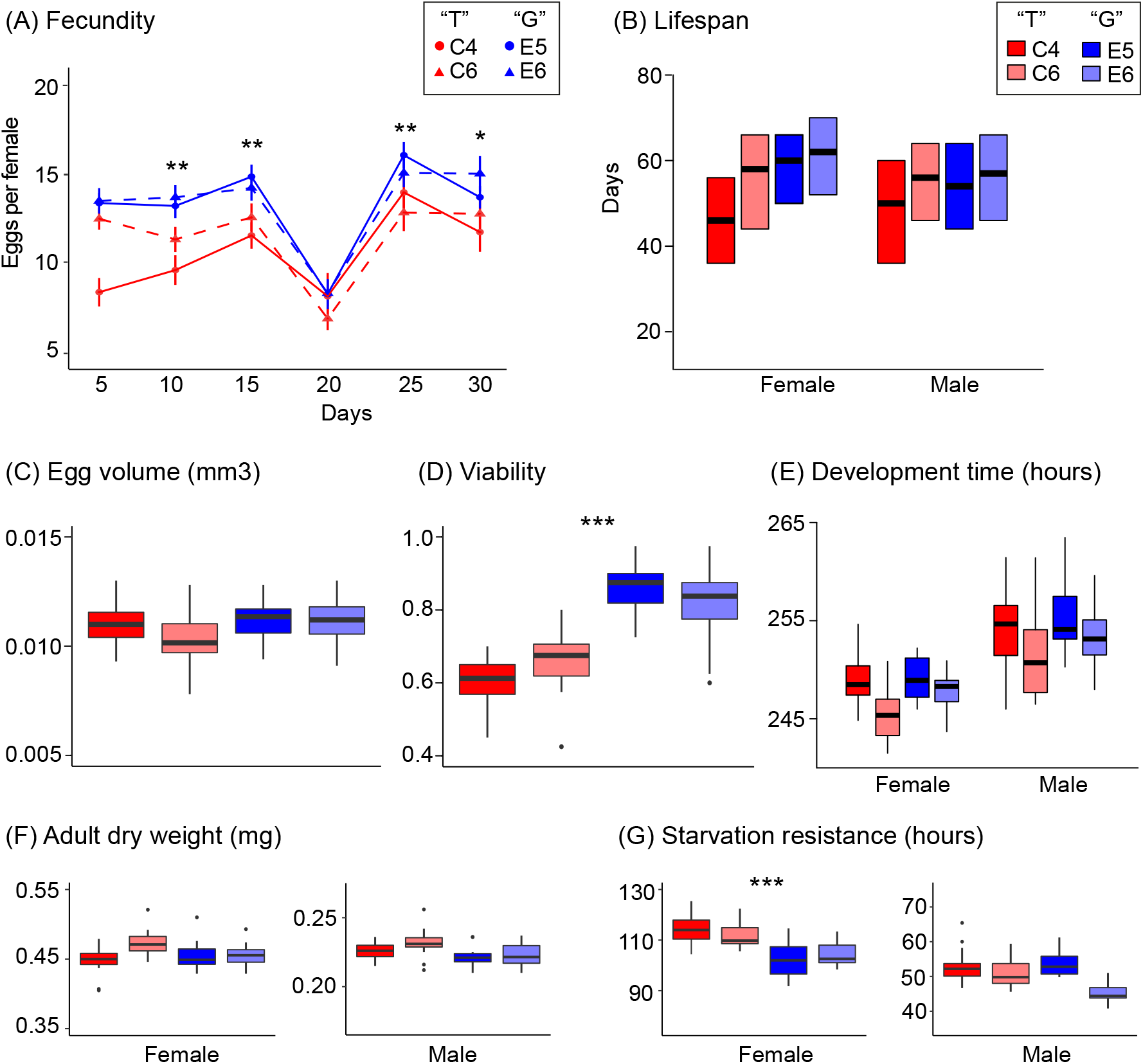
CRISPR/Cas9 modification of the *Eip75B* SNP in an isogenic background reveals its effects on life history. Two lines with the “G” allele (E5 and E6) and two lines with the “T” allele (C4 and C6) were constructed using CRISPR/Cas9 and tested for differences in a range of life history traits known to be regulated by *Eip75B* from mutant analyses and RNAi studies. (A) There was a significant effect of SNP identity on fecundity, as measured by daily egg-laying rates per female up to day 30 after emergence (allele: χ^2^ = 13.3, *P* = 0.0013, allele*day: χ^2^ = 3.8, *P* = 0.052), Also, there was a significant effect on (D) egg-to-adult viability (χ^2^ = 12.5, *P* = 0.0004), and (G) starvation resistance in females (females: χ^2^ = 11.9, *P* = 0.0006; males: χ^2^ = 0.7, *P* =0.41). This SNP allele did not have an effect on (B) lifespan (females: χ^2^ = 2.0, *P* = 0.16; males: χ^2^ = 1.4, *P* = 0.24), (C) egg volume (χ^2^ = 2.5, *P* = 0.12), (E) egg-to-adult development time (females: χ^2^ = 0.9, *P* = 0.0.34; males: χ^2^ = 1.7, *P* = 0.19), and (F) adult dry weight (females: χ^2^ = 0.3, *P* = 0.58; males: χ^2^ = 3.8, *P* = 0.052). Asterisks indicate significant results.

The significant effects on fecundity observed with CRISPR/Cas9 editing confirm the results of our association study; thus, two independent lines of evidence support the conclusion that the *Eip75B* SNP has a causal role in affecting this complex trait. To quantify the genetic contribution of this SNP to variation in daily egg-laying rates, we used the data from the association experiment to estimate the genetic variance attributable to the SNP and the proportional contribution of the SNP to the total variance (i.e., the intraclass correlation coefficient or “isofemale heritability”; see e.g., Hoffmann and Parsons 1988; David, et al. 2005) (Supplementary Table 2). These ‘back-of-the-envelope’ estimates suggest that this naturally segregating SNP at *Eip75B* might explain up to 13.5% of the variation in daily fecundity depending on the day.

The examined candidate SNP is located in an inferred cis-regulatory motif in close proximity to the transcription start site of isoform G of the gene, suggesting that it may play a role in regulating gene expression or gene splicing. Although different isoforms of *Eip75B* have been shown to have different functions, isoform G has not been studied yet. The precise molecular and physiological mode of action of this SNP thus remains to be characterized. The effect of this SNP on reproduction might be linked to the well-known function of *Eip75B* in oogenesis (Belles and Piulachs 2015; Ables, et al. 2016), but could also indicate that allelic variants at this locus modify nutrient availability or the metabolic processes required for egg production, which might be linked to the small, but significant effect of this SNP on starvation resistance. For example, *Eip75B* is known to play a role in the mating-induced remodeling of the gut, which prepares the female body for the increasing energy demands associated with egg laying (Reiff, et al. 2015; Carvalho-Santos and Ribeiro 2018; Zipper, et al. 2020).

Based on results from our previous experimental evolution study we had hypothesized that the *Eip75B* variant examined here might also affect lifespan and mediate the trade-off between early fecundity and longevity (Hoedjes, et al. 2019). However, this hypothesis was not supported by either of the two functional tests of this study, demonstrating that one needs to be careful when interpreting candidate SNPs identified in experimental evolution studies. In addition, other life-history traits known to be regulated by *Eip75B*, such as development time and adult size, were also unaffected by this SNP. The fact that lifespan was affected by RNAi silencing of the whole gene but not by the SNP is consistent with the notion that naturally segregating molecular polymorphisms have fewer pleiotropic functions than null alleles (amorphic mutations) of the same gene (Stern 2000, 2011).

### Conclusion

How individual nucleotide variants contribute to complex, quantitative traits is a largely unresolved question in evolutionary genetics. While it is clear that quantitative traits are highly polygenic, with many loci of small effect contributing to trait variation (the ‘infinitesimal model’; Rockman 2012; Yang et al. 2015; Boyle et al. 2017; Yengo et al. 2022; Barton 2022), surprisingly little is known about the effect sizes and properties of individual variants (Orr and Coyne 1992; Seehausen et al. 2014; Enbody et al. 2022). A small, but growing number of functional studies are beginning to reveal that natural variants can make large contributions to variation in fitness-related traits (Schmidt et al. 2008; Durmaz et al. 2019; Mokashi et al. 2021; Schluter et al. 2021; Enbody et al. 2022). Our study provides a rare example of functional characterization of an individual naturally segregating SNP, and demonstrates that it has causal pleiotropic effects on two complex fitness traits, viability and fecundity. These results indicate that, although life-history traits are highly polygenic, individual SNPs can have remarkably large phenotypic effects.

## MATERIALS AND METHODS

### RNAi

Transgenic RNAi was performed using the ubiquitously expressing, mifepristone (RU486)-inducible *daughterless* (*da*)-GeneSwitch(GS)-GAL4 (Tricoire, et al. 2009) (courtesy of Véronique Monnier, Paris) to drive the expression of *Eip75B* UAS-RNAi constructs during the adult stage only, and avoid adverse effects on development. In addition, the GeneSwitch system makes it possible to compare the effects of RNAi in a single transgenic genotype (i.e. application of the drug *versus* no application of the drug), providing a robust control with regard to genotype. Two independent UAS-RNAi constructs obtained from the Vienna *Drosophila* RNAi Center (VDRC) (#108399 [E10] and #44851 [E44]), and two additional independent constructs obtained from Bloomington *Drosophila* Stock Center (BDSC) (#TRiP.GLC01418 [E01] and #TRiP.JF02257 [E03]) were used. All constructs target a domain of *Eip75B* that is shared by all isoforms.

All lines were kept and assays were performed at 25°C, 65% humidity and 12h:12h light:dark cycle, on a cornmeal-yeast-sucrose-agar medium (per 1 liter of food: 7 g agar, 50 g sucrose, 50 g cornmeal, 50 g yeast, 6 ml propionic acid, 10 ml of a 20% nipagin stock solution). After emergence, flies were kept on medium containing either 100 or 200 μg/ml (233 or 466 μM, respectively) mifepristone dissolved in ethanol, or on control medium (i.e., ethanol without mifepristone). These concentrations have previously been used without detrimental effect on survival of adult flies (Osterwalder, et al. 2001; Tricoire, et al. 2009; Fabian, et al. 2018). To confirm this, we tested lifespan, fecundity and egg volume (as detailed below) of a cross of *da*-GS-GAL4 females with males of the isogenic host strain for the VDRC RNAi library, *w^1118^* (#60000) (Supplementary Figure 1).

Quantitative real-time PCR (qRT-PCR) was used to confirm knockdown efficiency of the different constructs. For this, cohorts of F1 offspring between *da*-GS-GAL4 females and males carrying one of the UAS-RNAi constructs were collected upon emergence, and placed in vials with either control medium or medium with mifepristone (100 or 200 μg/ml) for four days in groups of 10 females and 10 males. The flies were then flash frozen in liquid nitrogen and three females were pooled together per sample and homogenized for whole-body RNA extraction. RNA extractions were done using a MagMax robot, using the MagMax-96 total RNA isolation kit (ThermoFisher Scientific), following the instructions of the manufacturer. cDNA was prepared following the GoScript Reverse Transcriptase system (Promega); qRT-PCR was done on a QuantStudio 6 real-time PCR machine (ThermoFisher Scientific), using Power SYBR Green PCR Master mix (ThermoFisher Scientific). We analyzed ten biological samples per construct and mifepristone treatment, with three technical qRT-PCR replicates each. The expression levels of *Eip75B* and four reference genes (*TBP, Cyp1, Ef1a48D* and *Rap2l;* see Supplementary Table 3 for primer sequences) were measured. The reference genes had a stability of M<0.25 as determined *a priori* from a set of female wholebody samples of various strains and mifepristone concentrations using the GeNorm algorithm incorporated in qbasePLUS v.1.5 (Biogazelle). The three technical replicates were analyzed for reproducibility, and measurements were accepted if none or only one of the technical replicates differed >0.5 Cq from the other two replicates. In the latter case the divergent Cq value was removed from the dataset. All measurements fulfilled these criteria. The technical replicates were averaged and a single gene expression value per gene per sample was calculated using qbasePLUS v.1.5. The gene expression levels of the four reference genes were then averaged geometrically to obtain an accurate normalization factor for each sample (Vandesompele, et al. 2002). The relative gene expression levels of *Eip75B* for the four RNAi constructs were then calculated compared to the expression of the control treatment (i.e. no mifepristone) for each construct separately, respectively. The gene expression levels were analyzed in R (v.3.3.1) using ANOVA with mifepristone concentration as a fixed effect (continuous variable).

We detected a significant reduction in whole-body *Eip75B* expression three out of four constructs when being fed mifepristone (E01: *F* = 6.11, *P* = 0.020; E03: *F* = 19.66, *P* = 0.00013; E44: *F* = 4.51, *P* = 0.043) (Figure 1 and Supplementary Table 1). The average reduction in *Eip75B* expression was 18%, 28%, and 18%, respectively, at a mifepristone concentration of 200 μg/ml. No significant reduction in the expression of *Eip75B* was detected for construct E10 (*F* = 1.95, *P* = 0.04).

We examined the effects of *Eip75B* knockdown on lifespan, fecundity and egg volume. To measure the effects of lifespan, cohorts of F1 offspring between crosses of *da*-GS-GAL4 virgin females and males carrying one of the four UAS-RNAi constructs or the isogenic control strain were collected within a 24-hour window; the flies were sexed under mild CO_2_ exposure and transferred at random to 1-liter demography cages with food vials (with 0, 100, or 200 μg/ml mifepristone) attached to the cages. For each genotype and mifepristone concentration, we set up three replicate cages, each containing 75 flies per sex. Dead flies were scored and fresh food was provided every two days. Differences in lifespan between mifepristone-induced RNAi and uninduced controls were analyzed in R (v.3.3.1) for the two sexes separately using mixed-effects Cox (proportional hazards) regression with mifepristone concentration as fixed effect and with ‘replicate cage’ as a random effect using the R package *coxme* (v2.2-5) for each construct separately. In addition, to test the statistical significance of similar trends in lifespan changes across the four constructs, we used a mixed-effects Cox regression on all data, with mifepristone concentration, construct, and their interaction as fixed effects and with ‘replicate cage’ as random effect. Bonferroni correction was applied to account for multiple testing (0.05/8 = 0.00625).

The effect of *Eip75B* knockdown on fecundity was assessed by measuring daily egg-laying rates. For this, virgin females were collected from the F1 offspring of *da*-GS-GAL4 females and males from a UAS-RNAi strain or the isogenic control strain. After 24 hours, two virgin females and two *w^1118^* males were placed together in vials with 0 (i.e. control) or 100 μg/ml mifepristone. The analyses were limited to mifepristone concentrations of 100 μg/ml, as a significant effect of 200 μg/ml of mifepristone on the control was observed (Supplementary Figure 1). Ten replicate vials were prepared per genotype and mifepristone concentration. The flies were left for 48 hours to ensure mating and consumption of mifepristone before the start of the experiment. Then the flies were transferred to a fresh vial with food (with 0 or 100 mifepristone, respectively) to lay eggs for 24 hours. The number of eggs laid per pair of flies was counted under a microscope. The daily egg-laying rate was measured at day 3, 6, 9, 12 and 15.

After day 15, all flies were transferred to vials with medium without mifepristone for 24 hours. The eggs laid were collected and 30 randomly chosen eggs per strain and mifepristone concentration were photographed with a Canon EOS 60D camera under a Leica dissection microscope. The length and the width of each egg were measured using ImageJ (v.1.51h) software and the volume of each egg was approximated following the formula described by (Jha et al., 2015): volume = 1/6π(width)^2^*length. Variation in egg-laying rates was analyzed in R (v.3.3.1) using generalized linear mixed models with a Poisson distribution and with mifepristone concentration, day and their interaction as fixed effects and with ‘replicate vial’ as a random effect using the R package *lme4* (v.1.1-13). Egg volume was analyzed in R using ANOVA with mifepristone concentration as fixed effect. A Bonferroni correction was applied to account for multiple testing (0.05/4 = 0.0125).

### SNP association study

Strains from the *Drosophila* Genetic Reference Panel (DGRP) (Mackay, et al. 2012), obtained from the Bloomington *Drosophila* Stock Center (BDSC) were used for a SNP association experiment based on Mendelian randomization. The principle of this approach is that putative causal effects of loci are assessed by testing the alternative allelic states in a genetically diverse background to level out the effects of confounding epistatic factors. The SNP genotypes of our candidate SNP in *Eip75B* of each line were obtained from the genome database available for this panel. The genotype was confirmed by sequencing a small genomic region surrounding the SNP using Sanger sequencing. We generated ten independent, homozygous F1 crosses per allelic variant (i.e. “GG” or “TT”) from twenty DGRP strains each, and 18 independent F1 crosses that were heterozygous for the candidate SNP (i.e. “TG”), and then assessed lifespan and fecundity of these crosses. The F1 crosses were generated by crossing virgin females of one strain with males from another strain (see Supplementary Table 4). The F1 crosses were reared and assays were performed at 25°C, 65% humidity and 12h:12h light:dark cycle, on a cornmeal-yeast-sucrose-agar medium as described above.

Lifespan was assessed in demography cages as described earlier with the regular medium provided in food vials. Flies that emerged within a 24-hour window were collected and 75 males and 75 females were placed in a single demography cage. For the experiment using F1 crosses, a single demography cage per cross was setup. Differences in lifespan between SNP genotypes were analyzed in R (v.3.3.1) using mixed-effects Cox (proportional hazards) regression with allele, sex and their interaction as fixed effects and with ‘F1 cross’ as a random effect using the R package *coxme* (v2.2-5).

Egg-laying rate over a period of 30 days after emergence was measured to provide insight in both early (peak) and late (post-peak) fecundity. Again, flies that emerged within a 24-hour window were collected, and two females and two males were placed together in a vial containing regular medium. Three replicate vials per cross were setup. Every third day (i.e. day 3, 6, 9, 12, 15, 18, 21, 24, 27 and 30), the number of eggs laid per pair of flies during a period of 24 hours was counted under a microscope. Variation in egg-laying rates was analyzed in R (v.3.3.1) using generalized linear mixed models with a Poisson distribution and with SNP genotype, day and their interaction as fixed effects and with ‘replicate vial’ as a random effect using the R package *lme4* (v.1.1-13). Post-hoc Tukey tests were applied to test for significant differences between the three genotypes using the R package *emmeans*(v.1.7.0).

### CRISPR/Cas9-induced SNP modification

CRISPR/Cas9 makes it possible to edit genomes with the precision of a single nucleotide, but a relatively low efficiency of homology-directed repair in the fruit fly, combined with the necessity of molecular screening to detect a single-nucleotide modification, makes the procedure labor intensive (Ramaekers, et al. 2019; Zirin, et al. 2021). We therefore used a co-CRISPR or co-conversion approach, in which an unlinked (visible) marker for rapid screening is introduced simultaneously with the desired genomic modification, by injecting two different gRNAs and donor templates (Ge, et al. 2016; Kane, et al. 2017; Ewen-Campen and Perrimon 2018). The rationale underlying this approach is that the two modifications often co-occur resulting in an enrichment of the desired genomic modification in flies with the marker gene. Here we used DsRed as a visible marker to facilitate screening.

For editing the SNP in *Eip75B* at position 3L:18026199, plasmids containing the gRNA sequence and dsDNA donor templates were prepared and injected into *y,w;nos-Cas9* (II-attP40)/*CyO* embryos (strain obtained from BestGene Inc. (Chino Hills, CA, USA); it was generated by backcrossing the original strain NIG-FLY CAS-0001 into a *y,w* background). This strain has a “T” allele at position 3L:18026199; hence the aim of the experiment was to modify it to a “G” allele. In addition to editing the candidate *Eip75B* SNP, a DsRed marker was inserted to chromosome 2L, into the C-terminus of *I(2)gl*, to facilitate screening. For this, gRNA sequences were cloned into bicistronic pDCC6 plasmids (*Eip75B:* 5’ CACTGCTTGCAATAAGTGATGGG 3’;*I(2)gl*: 5’ TGTTAAAATTGGCTTTCTTCAGG 3’). Homology directed repair was facilitated by constructing donor templates matching the flanking sequences of the injection line for 500bp on either side of the dsDNA cleavage point (i.e. 1 kb in total). These donor templates were cloned into pcDNA3.1. The injection mix consisted of 35 μg of the *Eip75B* donor plasmid, 35 μg of the *I(2)gl* dsRED donor plasmid, 15 μg I2gl gRNA plasmid, and 15 μg *Eip75B* plasmid per 100μl. The CRISPR/Cas9 editing was done by InDroso (Rennes, France). A total of 300 embryos were injected, resulting in 60 surviving larvae and 30 fertile adults. The flies were crossed to a balancer line (*w;+;Dr/TM6,Sb,Tb;* BDSC #8576) to facilitate the generation of homozygous lines (see Supplementary File 3 for the complete crossing scheme used and a discussion on genetic background). The offspring of the cross was screened visually for the DsRed marker and molecularly, by sequencing, for the *Eip75B* candidate SNP. Lines that were positive for the DsRed marker, but negative for the modification in *Eip75B* (hence, they carried the “T” allele) were kept to serve as controls. The *Eip75B*-modified and control lines were backcrossed to the balancer line for three generations to remove the DsRed marker and to obtain homozygous lines for the candidate SNP: *w;;Eip75B^T18’026’199G^* (edited) and *w;;Eip75B*^T^ (control). We generated two homozygous edited lines with the “G” allele (“E5” and “E6”) and two lines with the “T” allele (“C4” and “C6”). The sequence surrounding the *Eip75B* SNP, including the full length of the homology arms, were sequenced to confirm the identity of the SNP and to monitor for any potential undesired genomic modifications. No genomic modifications, apart for the candidate SNP, were detected in any of the lines (Supplementary File 3).

The following phenotypes were tested in the CRISPR-edited and control lines: lifespan, egg-laying rate, egg volume, egg-to-adult development time, egg-to-adult viability, starvation resistance, and adult dry weight. Before the onset of phenotyping experiments flies that were used to generate the cohorts for the experiments were also screened for the identity of the *Eip75B* SNP using PCR-based screening. For this, gDNA was isolated from individual flies using Chelex extraction (Biorad). Flies were placed individually in 100 ul Chelex solution (5g of Chelex mixed with 95ml TE buffer) with 5 ul proteinase K (Promega), and incubated overnight at 56°C. The resulting gDNA was then analyzed with two alternative reverse primers to distinguish the two allelic states: forward primer 5’ CCGGGCACAGTCTCAGTGTT 3’, in combination with reverse primers 5’ GCGCCAAAGCTTTCCCGTC**<u>A</u>**3’ or 5’ GCGCCAAAGCTTTCCCGTC**<u>C</u>**3’. The PCR was done with GoTaq2 polymerase following the instructions of the manufacturer (Promega) and the following PCR protocol: 94°C for 3 min; 32 cycles with 94°C for 15 sec, 62°C for 30 sec, and 68°C for 1 min; 68°C for 6 min. This procedure facilitated rapid and high-throughput screening by gel electrophoresis. A subset of the samples was sequenced using Sanger sequencing for confirmation. All phenotyping experiments were done in parallel using the same cohort of flies.

Lifespan was measured as described above, by placing 75 females and 75 males that emerged in a 24h window in demography cages with standard medium provided. Five replicate cages were set-up per strain. Differences in lifespan between the two allelic states in *Eip75B* were analyzed in R (v.3.3.1) using mixed-effects Cox (proportional hazards) regression with allele, sex and their interaction as fixed effects and with ‘line’ and ‘replicate cage’ as random effects using the R package *coxme* (v2.2-16).

Egg-laying rate was measured as described above. Two females and two male flies, emerged within a 24-hour window, were placed together in a vial containing standard medium. On days 5, 10, 15, 20, 25, and 30 the number of eggs laid per pair of flies during a period of 24 hours was counted under a microscope. Egg-laying rates was analyzed in R (v.3.3.1) using generalized linear mixed models with a Poisson distribution and with allelic state, day and their interaction as fixed effects and with ‘line’ and ‘replicate vial’ as random effects using the *R* package *lme4* (v.1.1-26). The effect of allelic state for each day separately were analyzed in models with only allelic state as fixed effect and ‘replicate vial’ as random effect.

Egg volume was assessed by collecting eggs from females, age 7 days after emergence, laid overnight. We randomly picked 30 eggs, which were photographed and measured as described above. Egg volume was analyzed in *R* using ANOVA with allele as fixed effect.

To assay egg-to-adult development time ~400 flies were placed in bottles with regular medium and yeast added to lay eggs during two consecutive periods of 3 hours (“batch 1” and “batch 2”). Eggs were collected and distributed to vials at density of 40 eggs per vial, with 16 replicate vials per line. Emergence of adult flies was scored three times per day, at 08:00 (“lights on”), 14:00, and 20:00 (“lights off”). For each vial, the mean time of emergence per sex per vial was calculated and analyzed using linear mixed models with allele, sex and their interaction as fixed effects and batch and line as random effects using the *R* package *lme4*. The egg-to-adult viability was calculated as the proportion of flies that emerged from 40 eggs per vial. It was analyzed using linear mixed model with allele as fixed effect and batch and line as random effects using *lme4*.

Starvation resistance was measured by collecting flies from a 24h window of emergence and placing them in vials with standard medium at a density of 10 females and 10 males for 5 days. Then the two sexes were placed in separate vials. At day 7 after emergence, the flies were transferred to vials with 0.5% agar. Survival was scored three times daily (08:00 (“lights on”), 14:00, and 20:00 (“lights off”)) until all flies has died. The mean time to death was calculated per vial and analyzed for both sexes separately using linear mixed models with allele as fixed effect and line as random effect.

Adult dry weight was assayed by collecting flies from a 24h window of emergence and placing them in vials with standard medium at a density of 10 females and 10 males. On day 7 after emergence, the flies were frozen at −20°C. Flies were dried individually by placing them in 96-wells plates in an incubator at 65°C for three days. This was done in two batches. Adult dry weight was then determined in pools of six flies, with 16 pools of flies per line and sex, on a Mettler Toledo MT5 microbalance. The mean adult dry weight per fly was analyzed for both sexes separately using linear mixed models with allele as fixed effect and line as random effect.

### Genetic contribution of the SNP to fecundity

After demonstrating a causal effect of the *Eip75B* SNP (3L:18026199) on fecundity, we sought to estimate the contribution of this polymorphism to fecundity using the data from the F1 SNP association study. To this end, we first calculated the population mean (*M;* mean genotypic value) using the average effects of the three genotypes (i.e., their genotypic values; the mean number of eggs laid per female) and the allele frequencies (e.g., see (Falconer and Mackay 1996; Lynch and Walsh 1998) as follows:

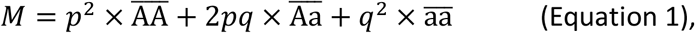

where 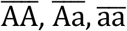 represent the genotypic values of “TT”, “TG”, and “GG”, respectively, and *p* and *q* are the population frequencies of the *A* (=“T”) and *a* (=“G”) alleles, respectively. For *p* and *q* we used the allele frequencies observed in the E&R study (Hoedjes, et al. 2019): *p* = 0.68 (“T”) and *q* = 0.32 (“G”) (frequencies averaged across the experimentally evolved replicate populations).

By re-expressing the effects in Equation 1 as deviations from the population mean, we estimated the genetic variance due to the SNP as:

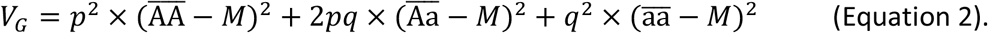

However, the F1 crosses do not faithfully represent the 3 genotypic classes and their frequencies in the experimental evolution experiment; and *p*^2^, 2*pq* and *q*^2^ represent the expected, not the observed genotype frequencies, estimated from the observed allele frequencies. As an alternative approach, instead of using Equation 1 based on allele frequencies, we therefore also estimated *V_G_* by calculating the variance among the three genotypes with a random effects model (general linear mixed model with “Genotype” as random effect) in R (v.3.3.1) using the *lme4* (v.1.1-13) package. Both approaches yielded qualitatively similar variance estimates (Supplementary Table 2).

Based on these variance estimates, we quantified the proportion of the total variance attributable to the SNP, i.e., the intraclass correlation coefficient *t* (also called “isofemale heritability”; e.g., see (Hoffmann and Parsons 1988; Falconer and Mackay 1996; David, et al. 2005), as:

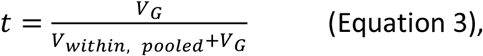

where *V_within, pooled_* represents the variance among line means within each genotype, pooled across all genotypes:

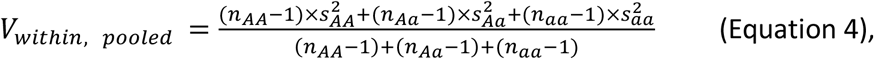

where *n* represents the sample sizes and *s^2^* the variances of the 3 genotypes. The intraclass correlation coefficients reported in Supplementary Table 2 thus provide a quantification of the contribution of the *Eip75B* variant to the trait of interest; this being said, the above described estimations only represent “back-of-the-envelope” calculations.

## ACKNOWLEDGEMENTS

We thank Tadeusz Kawecki for generously allowing us to use his *Drosophila* facilities; the Flatt and Keller labs for discussion and support; Véronique Monnier for the *da*-GS-GAL4 strain; and the Bloomington *Drosophila* Stock Center (BDSC) and the Vienna *Drosophila* RNAi Center (VDRC) for providing stocks. Our work was supported by H2020-MSCA-IF-2015 (grant 701949 to KMH), the European Research Council (grant 741491 to LK), and the Swiss National Science Foundation (grants PP00P3_165836, PP00P3_133641/1, and 310030E-164207 to TF, grant 310030B_176406 to LK).

## DATA AVAILABILITY

All raw data will be made publicly available on Dryad upon acceptance of the manuscript.

