## Supplementary data for "A single nucleotide variant in the PPARγ-homolog *Eip75B* affects fecundity in *Drosophila*"

Supplementary File 1: ***Population genetic analysis of the candidate SNP in Eip75B***

Supplementary File 2: ***Linkage disequilibrium within Eip75B***

Supplementary File 3: ***CRISPR/Cas9 approach and subsequent crossing scheme***

Supplementary Figure 1: ***Control experiment for adverse phenotypic effects of mifepristone***

Supplementary Table 1: ***Confirmation of knockdown of Eip75B by RNAi***

Supplementary Table 2: ***Estimation of genetic contribution of the Eip75B SNP to fecundity***

Supplementary Table 3: ***qRT-PCR primers used to quantify Eip75B knockdown***

Supplementary Table 4: ***F1 crosses for SNP association study***

**Supplementary File 1**

***Population genetic analysis of the candidate SNP in Eip75B***

***Methodology***

The frequency of the "T" allele of the candidate SNP in *Eip75B* (3L:18026199, release 6) ranged from an average of 0.52 to 0.84 in early reproducing *versus* late reproducing, long-lived experimentally evolved (EE) populations, respectively (Hoedjes, et al. 2019). These allele frequencies were compared to the allelic variation that exists in populations from Europe, North-America, and Africa similar to (Hoedjes, et al. 2022). For this, we analyzed whole genome sequencing data from multiple geographic populations available from the DEST dataset (Kapun, et al. 2021). We included both single individual sequencing data and Pool-Seq data from various sources. For the single individual sequencing data, the "T" allele frequency of the *Eip75B* allele was calculated when at least ten individuals were sequenced for a given population. Data from pooled populations was included when the coverage for each *Eip75B* SNP was at least 10. We tested the data from the 173 samples collected in Europe for clinal distribution patterns of the allele frequencies. For this, we used generalized linear models (GLMs) based on the allele counts of the *Eip75B* SNP, with either longitude or latitude as explanatory factor. False discovery rate was estimated by testing clinal distribution patterns among a set of 21'008 neutral SNPs (Parsch, et al. 2010; Clemente and Vogl 2012). Empirical cumulative density functions (ECDF) were generated based on the *P* values of these neutral SNPs. The area of the upper tail confined by the $\chi2$ values of the *Eip75B* SNP indicates the percentile of neutral SNPs with $\chi2$ values equal or higher than the *Eip75B* SNP. This parameter provides a corrected estimate of significance of clinality of each SNP compared to genome-wide neutral estimates (Ramaekers, et al. 2019).

***Distribution of the Eip75B allele***

Analysis of the allele frequencies of the *Eip75B* SNP in multiple geographic populations from Europe (173 samples, "T" allele frequency: 0.53-0.92) and North America (55 samples: 0.53-0.82) indicates a variable allele frequency for this SNP in natural populations, and demonstrates that the allele frequencies observed by (Hoedjes, et al. 2019) fall within the range of naturally occurring variation in Europe and North America (Supplementary Table 1). Limited data on the allele frequencies of the candidate *Eip75B* SNP in African populations suggests that both alleles were present in ancestral populations already, although the frequency of the "G" allele may have been limited ("T" allele frequency: 0.8 in Gabon, 1.0 in Cameroon, 0.99 in Zambia). Analysis of the allele frequency data from the European populations provided no indication for a clinal distribution of the *Eip75B* SNP (latitude: $\chi^2$ = 1.13, $P$ = 0.29, $P_{\text{corrected}}$ = 0.61; longitude: $\chi^2$ = 9.45, $P$ = 0.002, $P_{\text{corrected}}$ = 0.43).

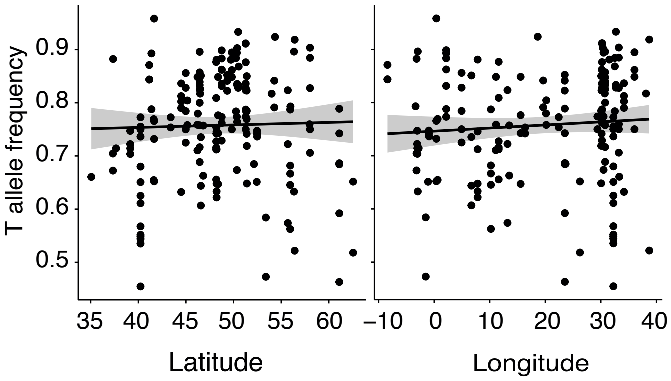


**Supplementary File 1 – figure 1:** Allele frequency distribution of the "T" allele at position 3L: 18026199 across Europe. The samples are ordered by latitude (left) and longitude (right).

**Supplementary File 2**

### *Linkage disequilibrium within Eip75B*

To estimate the potentially confounding effects of SNPs that are located closely to our candidate *Eip75B* SNP (3L:28'026'199) we estimated LD (measured by pairwise $r^2$) of all SNP (that have a minor allele frequency ≥ 0.1) located within *Eip75B* in the full panel of DGRP lines, following the approach previously published by as done by (Durmaz, et al. 2019; Hoedjes, et al. 2022). These analyses indicate very low levels of LD across the entire gene (Figure 1). This observation agrees with previous observations that LD decays very rapidly, within a few hundred base pairs, in *D. melanogaster* (Mackay, et al. 2012). The Mendelian randomization approach was, therefore, expected to provide information on the functional impact of the *Eip75B* SNP, with little or no confounding effects of the genetic background.

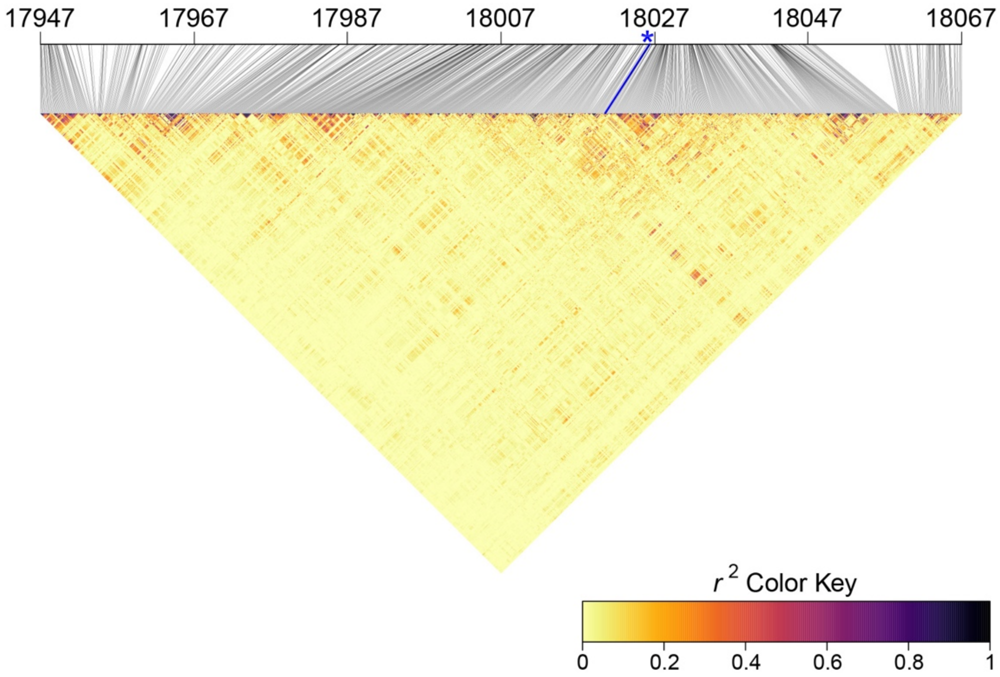


**Supplementary File 2 - figure 1:** Linkage disequilibrium (measured by pairwise $r^2$) of SNPs within *Eip75B*. The candidate SNP is indicated with a blue asterisk; the coordinates are provided in kb.

**Supplementary File 3**

***CRISPR/Cas9 approach and subsequent crossing scheme to generate edited ("G") and control ("T") lines***

**Injection of gRNA and dsDNA donor into embryos**

We used a transgenic *D. melanogaster* line that expressed the Cas9 construct under the control of a germline-specific *nos* promotor (*yw*;*nos*-Cas9/*CyO*) in order to induce the formation of heritable genome modifications. The transgenic *nos*-Cas9 construct is located on chromosome 2; it originated from the line NIG-FLY #CAS-0001, and was backcrossed into a *yw* background to improve line health by BestGene Inc (Chino Hills, CA, USA). A single batch of embryos (N=300) from the injection line was injected with the gRNA and the dsDNA donor construct as detailed in the Methods section, with the aim of generating flies with the modified SNP and a visible DsRed marker in the DNA of their germline. These flies were subjected to a crossing scheme consisting of 7 crosses to (1) be able to track chromosome 3, on which the target *Eip75B* SNP is located, for molecular screening, (2) to make chromosome 3 homozygous, and (3) to replace chromosome 2 of the injection line, which has the *nos*-Cas9 transgene, with a wildtype ("+") chromosome.

**Cross 1**

The injections yielded 60 surviving larvae, resulting in 30 fertile adults. Each adult was collected as a virgin and was then mated individually to the balancer line (*w*;+; *Dr*/TM6B,*Sb*,*Tb*; BDSC #8576).

**Cross 1:**

| *1 male (injection line)* | x | *1 female (balancer line)* |
|---|---|---|
| $yw;\ \frac{nos\ \mathrm{Cas9\ (attP40)},I(2)gl\ \mathrm{DsRed}}{CyO};\ Eip75B^{\mathrm{T\ or\ (}}$ | x | $w;+;\ \frac{Dr}{\mathrm{TM6B},Tb,Sb}$ |

The 30 resulting pools of offspring were inspected for the presence of the visible DsRed marker, which was observed in 7 pools of (F1) offspring. These pools were considered the “jackpot” pools, with a higher chance of also harboring the desired SNP modification (i.e., the principle of co-CRISPR). These pools of offspring typically contain a mixture of the original and the edited genotypes as CRISPR/Cas9 resulted in a mosaic of transformed and non-transformed germline cells in the injected individuals, necessitating molecular screening of individual flies for the identity of the *Eip75B* SNP. Selected offspring from these pools were used to set up the second cross and for molecular screening to assess the target SNP identity. Note that the four lines (“E5”, “E6”, “C4”, and “C6”) all originated from different pools of offspring resulting from cross 1 (i.e., the offspring of different injected individuals), hence the lines represent independent replicates (see figure 1).

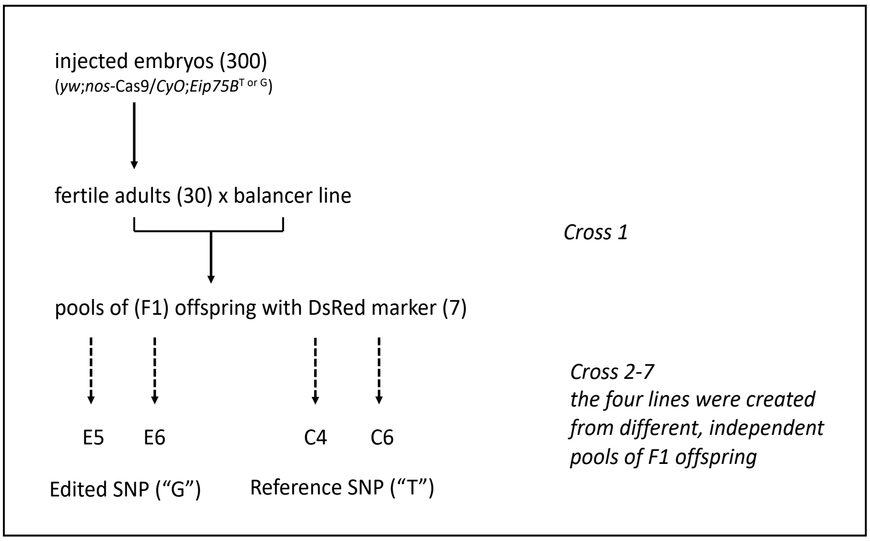


**Supplementary File 3 - figure 1:** Overview of the crossing scheme and establishment of 4 independent CRISPR/Cas9 lines ("E5", "E6", "C4", and "C6"). The batch of 300 injected embryos yielded 30 fertile adults. These adults were collected as virgins and subsequently mated individually to one fly (of the opposite sex) of the balancer line. Pools of offspring from each of these 30 pairs were observed for the expression of the DsRed marker. The lines "E5", "E6", "C4" and "C6" have been derived from different F1 pools (and hence, ultimately, from different injected individuals) and have gone through the exact same crossing scheme, completely independent from each other.

<u>**Cross 2**</u>

Offspring from cross 1 that expressed the DsRed marker were backcrossed individually and screened for the presence of the "G" allele (eventually resulting in edited lines "E5" and "E6") (after crosses 2-7). Flies that expressed the DsRed marker but carried the "T" allele were also backcrossed to construct the control lines ("C4" and "C6"). The four CRISPR lines ("E5", "E6", "C4", and "C6") were derived from different pools of F1 offspring and, hence, from different, independently injected embryos. All four lines followed the exact same crossing scheme, independent from each other.

**Cross 2:**

| *1 male (selected offspring of cross 1)* | x | *1 female (balancer line)* |
|---|---|---|
| $w; \frac{+}{CyO}; \frac{Eip75B^{\mathrm{T\ or\ G}}}{\mathrm{Tm6B}, Tb, Sb}$ | x | $w; +; \frac{Dr}{\mathrm{TM6B}, Tb, Sb}$ |

**Cross 3: Established balanced line**

Offspring of cross 2 with the desired *Eip75B* SNP ("T" or "G"), and with the *nos*-Cas9 construct and the *CyO* balancer removed from the second chromosome, were mated to each other to generate established balanced stock lines. For this, 5 males and 5 females with the desired genotype from a single pool of offspring were used. The exact same cross was done for each of the four independent CRISPR lines.

**Cross 3:**

| *5 males (selected offspring of cross 2)* | x | *5 females (selected offspring of cross 2)* |
|---|---|---|
| $w; +; \frac{Eip75B^{\mathrm{T\ or\ G}}}{\mathrm{Tm6B}, Tb, Sb}$ | x | $w; +; \frac{Eip75B^{\mathrm{T\ or\ G}}}{\mathrm{TM6B}, Tb, Sb}$ |

**Cross 4: Established homozygous line**

Offspring of cross 3 that were homozygous for the desired *Eip75B* (i.e., without the TM6B balancer chromosome) were mated to each other to generate homozygous balanced stock lines. For this, 5 males and 5 females with the desired phenotype/genotype from a single pool of offspring were used. A region of approximately 1 kb surrounding the SNP was sequenced to confirm the desired SNP and a lack of any off-target effects in the region (see below). The exact same cross was done for each of the four independent CRISPR lines. Generating the homozygous lines, which are the result of this cross, will make it easier maintain the induced SNP modification, as it excludes the necessity of screening the flies every generation. These established homozygous lines could be screened for effects on the phenotype of interest.

Cross 4:

| *5 males (selected offspring of cross 3)* | x | *5 females (selected offspring of cross 3)* |
|---|---|---|
| $w;+;\frac{Eip75B^{\mathrm{T\ or\ G}}}{Eip75B^{\mathrm{T\ or\ G}}}$ | x | $w;+;\frac{Eip75B^{\mathrm{T\ or\ G}}}{Eip75B^{\mathrm{T\ or\ G}}}$ |

**<u>Crosses 5-7</u>**

In our experiment, we opted to add three additional crosses to our crossing scheme, after generating the established homozygous lines (cross 4). This was done as a precaution as it took several months to generate and evaluate the genotypes of all four homozygous lines, and because the lines were shipped during this period. All four lines were simultaneously submitted to a second round of backcrossing, using the same TM6B balancer line as in the earlier crosses, prior to starting the phenotypic experiments. The 3 additional crosses served multiple purposes: (1) to be able to check the genotypes of the lines once more by molecular screening and to sequence the area surrounding the SNP; (2) to homogenize the genomic background of the lines, in particular the background of chromosomes 2 and X, as much as possible; and (3) to counter the (potential) accumulation of mutations in the genomic background over time. Immediately after these crosses, the lines were amplified and used in phenotyping assays. In cross 5, 5 male flies of the homozygous CRISPR/Cas9 lines were each mated individually to 3 females of the balancer line and genotyped to confirm their SNP identity.

**Cross 5:**

| *5x1 males (individuals of established homozygous stock; i.e. offspring of cross 4)* | x | *5x3 females (balancer line)* |
|---|---|---|
| $w; +; \frac{Eip75B^{\mathrm{T\ or\ G}}}{Eip75B^{\mathrm{T\ or\ G}}}$ | x | $w; +; \frac{Dr}{\mathrm{TM6B}, Tb, Sb}$ |

Cross 6 established balanced stocks by mating 10 male and 10 female offspring from each of the 5 pools of offspring of cross 5 with the desired genotype. Cross 7 established the final homozygous stock by mating 10 male and 10 female offspring from each of the 5 pools of offspring of cross 6 with the desired genotype.

**Cross 6: new balanced stock**

| | | |
|---|---|---|
| *10 males (selected offspring from each of the five pools of offspring of cross 5)* | x | *10 females (selected offspring from each of the five pools of offspring of cross 5)* |
| $w; +; \frac{Eip75B^{\mathrm{T\ or\ G}}}{\mathrm{Tm6B}, Tb, Sb}$ | x | $w; +; \frac{Eip75B^{\mathrm{T\ or\ G}}}{\mathrm{TM6B}, Tb, Sb}$ |

**Cross 7: final homozygous stock**

| | | |
|---|---|---|
| *10 males (selected offspring from each of the five pools of offspring of cross 6)* | x | *10 females (selected offspring from each of the five pools of offspring of cross 6)* |
| $w; +; \frac{Eip75B^{\mathrm{T\ or\ G}}}{Eip75B^{\mathrm{T\ or\ G}}}$ | x | $w; +; \frac{Eip75B^{\mathrm{T\ or\ G}}}{Eip75B^{\mathrm{T\ or\ G}}}$ |

After cross 7, the 5 pools of offspring (i.e., descending from the 5 males of the homozygous CRISPR lines and the 15 balancer females used in cross 5) were merged to generate the final lines (also see figure 2). This was done to prevent fixation of potentially confounding allelic variants on chromosomes 2 and X (e.g., due to drift), which might confound our ability to detect an effect of the focal SNP in *Eip75B*.

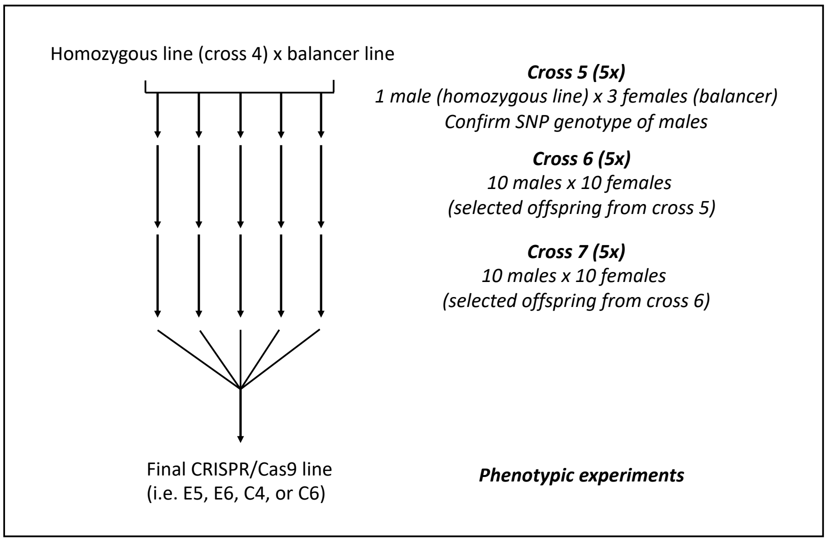


**Supplementary File 3 - figure 2:** Overview of the crossing scheme for crosses 5-7, and the generation of the final CRISPR/Cas9 lines

### Genetic variation and genomic background of the CRISPR/Cas9 lines

Our CRISPR/Cas9 lines were constructed using inbred laboratory strains (i.e., the injection line *yw*;*nos*-Cas9/*CyO* and the balancer line *w*;+;*Dr*/*TM6B,Tb,Sb*). Although these lines are expected to be inbred, residual heterozygosity can nonetheless be a confounding factor if (1) variants in the genetic background are in strong or complete linkage disequilibrium (LD) with the candidate SNP; and (2) if these linked background SNPs have an effect on the phenotype of interest. To investigate the effect of potentially confounding variants in the genomic background of our lines, we sequenced an area of ~1kb surrounding the edited *Eip75B* SNP of 5-10 individuals per line (sequences given below). We detected no residual heterozygosity in this region, i.e., all the sequenced individuals from the four lines had identical sequence, apart from the edited SNP. Given that LD is known to decay very quickly in *D. melanogaster*, typically within a few hundred base pairs (Mackay et al. 2012), it is therefore very unlikely that long-range LD confounds our results. In addition, we aimed to further limit any potential impact of long-range LD by homogenizing the genetic background (i.e., via crosses 5-7, homogenizing chromosomes 2 and X) and by using two independent CRISPR/Cas9 replicate lines for each allelic state (“G” versus “T”).

**<u>Final sequences of the region surrounding the target SNP</u>**

>Line E5

GCTCTGCTTTTGCTGCTGCTTCTGCTTTTTTGGGCCTTCGTAAGGCAGGACTAGGTTTTTGGTGCTTCTGTTGCC
AACCGGAGCGGTAATCAGACTTCGGCCGAGGATTTTATCCGCTGCATTCCACAAATCGCTTGTTGGTTTTAGC
GGCCACAATCTGATAGTTTCACATTTGGTTAAATGTTCGTCATCAGCCAATCACTTTTGTCTGGTCCGCCAATG
GGTTTCAGCAGAAAGGGTTAGGTCATTTAATTTTATTGGAACTGAAATATACACAAACTTATGTAAAGAAAGA
TTTAACTTTTCTACTTGATTAAACTACTGATAATGTTTAAACTGAATTCTTTAAATACAAAATTCTATACAACAAA
TTCTATAGAAACGAATTTCGCTTACAATGCACATAGGTCTAGTAAATAACACTTTGACTACAAAGGCAAATTTC
TAAAAGAAAATGTACTAATTTCCTATACTTAAACTAAACCTGTACTTAAACTAGTGGCACTGAGATTCAAAACC
AGGCGCCAAAGCTTTCCCATCCCTTATTGCAAGCAGTGCAAAGCTCCATGCAAGCAGTGCGACAAACTGTAGC
CACATGGCCGGCCCCAAAAAAAAAAGCAAAAAACAAAGCGAACAGCTGACTGAAACTTGCGATTTTCTGCTTCT
GCATTTCGGCTTTGGCGCCCCGCGTTTGTTGTTTTGCTTATTATTCTGCGTTCTTCGCTCGCTGCGCGAATACCG
AATTCTGAACACTGAGACTGTGCCCGGCGCTCAAAAATCTTTCCCAAACTTTTCGGCATTTCCACGTGAATTTG
AACGGCCAAAGCACAAATCAAAAACATATTTTTTGAGAATTACCCGAGTGTCATAAATAGAGAGAGTGATAA
GTCTTTCTAAATGTTTAATAATTAGAGAATGCATAAACAAATGGTGGCATTTATTAGATGAAAATGCATAAAG
ACTTGGCCAAAAATAAAACACGTAAAAAATCCTACATCATCTCTACTCTGGTTGCTACCTTGTGATAAAACTGT
GCACACAATTATTCCATAGGAAATTTAATCAAAGTTTTTTAAAGTTAAAGTTAAAAACAAGTTGATAGCCAAAA
CTAGAATAGTTAGCTAAACGAATTGAACACTCACAAAGGTGCCATAAAAAGCCTTAAATAGATTAACAAGA

>Line E6

GCTCTGCTTTTGCTGCTGCTTCTGCTTTTTTGGGCCTTCGTAAGGCAGGACTAGGTTTTTGGTGCTTCTGTTGCC
AACCGGAGCGGTAATCAGACTTCGGCCGAGGATTTTATCCGCTGCATTCCACAAATCGCTTGTTGGTTTTAGC
GGCCACAATCTGATAGTTTCACATTTGGTTAAATGTTCGTCATCAGCCAATCACTTTTGTCTGGTCCGCCAATG
GGTTTCAGCAGAAAGGGTTAGGTCATTTAATTTTATTGGAACTGAAATATACACAAACTTATGTAAAGAAAGA
TTTAACTTTTCTACTTGATTAAACTACTGATAATGTTTAAACTGAATTCTTTAAATACAAAATTCTATACAACAAA
TTCTATAGAAACGAATTTCGCTTACAATGCACATAGGTCTAGTAAATAACACTTTGACTACAAAGGCAAATTTC
TAAAAGAAAATGTACTAATTTCCTATACTTAAACTAAACCTGTACTTAAACTAGTGGCACTGAGATTCAAAACC
AGGCGCCAAAGCTTTCCCATCCCTTATTGCAAGCAGTGCAAAGCTCCATGCAAGCAGTGCGACAAACTGTAGC

CACATGGCCGGCCCCAAAAAAAAAGCAAAAAACAAAGCGAACAGCTGACTGAAACTTGCGATTTTCTGCTTCT
GCATTTCGGCTTTGGCGCCCCGCGTTTGTTGTTTTGCTTATTATTCTGCGTTCTTCGCTCGCTGCGCGAATACCG
AATTCTGAACACTGAGACTGTGCCCGGCGCTCAAAAATCTTTCCCAAACTTTTCGGCATTTCCACGTGAATTTG
AACGGCCAAAGCACAAATCAAAAACATATTTTTTGAGAATTACCCGAGTGTCATAAATAGAGAGAGTGATAA
GTCTTTCTAAATGTTTAATAATTAGAGAATGCATAAACAAATGGTGGCATTTATTAGATGAAAATGCATAAAG
ACTTGGCCAAAAATAAAACACGTAAAAAATCCTACATCATCTCTACTCTGGTTGCTACCTTGTGATAAAACTGT
GCACACAATTATTCCATAGGAAATTTAATCAAAGTTTTTTAAAGTTAAAGTTAAAAACAAGTTGATAGCCAAAA
CTAGAATAGTTAGCTAAACGAATTGAACACTCACAAAGGTGCCATAAAAAGCCTTAAATAGATTAACAAGA

>Line C4
CTGTTATTGGTGTTATTGCTATTGCTACTGGCCAACGCCGAAGCCTTTCTCACGCCTCGTTTTTGGCTCGTATCG
CTGTTCGTCAGCCCGGCCTTGAATAATTCTTCTGTCATTGGCATTTTCACATTTCATTTAACTTTGGCCCGCTCTG
CTTTTGCTGCTGCTTCTGCTTTTTTGGGCCTTCGTAAGGCAGGACTAGGTTTTTGGTGCTTCTGTTGCCAACCGG
AGCGGTAATCAGACTTCGGCCGAGGATTTTATCCGCTGCATTCCACAAATCGCTTGTTGGTTTTAGCGGCCACA
ATCTGATAGTTTCACATTTGGTTAAATGTTCGTCATCAGCCAATCACTTTTGTCTGGTCCGCCAATGGGTTTCAG
CAGAAAGGGTTAGGTCATTTAATTTTATTGGAACTGAAATATACACAAACTTATGTAAAGAAAGATTTAACTTT
TCTACTTGATTAAACTACTGATAATGTTTAAACTGAATTCTTTAAATACAAAATTCTATACAACAAATTCTATAG
AAACGAATTTCGCTTACAATGCACATAGGTCTAGTAAATAACACTTTGACTACAAAGGCAAATTTCTAAAAGA
AAATGTACTAATTTCCTATACTTAAACTAAACCTGTACTTAAACTAGTGGCACTGAGATTCAAAACCAGGCGCC
AAAGCTTTCCCATCACTTATTGCAAGCAGTGCAAAGCTCCATGCAAGCAGTGCGACAAACTGTAGCCACATGG
CCGGCCCCAAAAAAAAAGCAAAAAACAAAGCGAACAGCTGACTGAAACTTGCGATTTTCTGCTTCTGCATTTC
GGCTTTGGCGCCCCGCGTTTGTTGTTTTGCTTATTATTCTGCGTTCTTCGCTCGCTGCGCGAATACCGAATTCTG
AACACTGAGACTGTGCCCGGCGCTCAAAAATCTTTCCCAAACTTTTCGGCATTTCCACGTGAATTTGAACGGCC
AAAGCACAAATCAAAAACATATTTTTTGAGAATTACCCGAGTGTCATAAATAGAGAGAGTGATAAGTCTTTCT
AAATGTTTAATAATTAGAGAATGCATAAACAAATGGTGGCATTTATTAGATGAAAATGCATAAAGACTTGGCC
AAAAATAAAACACGTAAAAAATCCTACATCATCTCTACTCTGGTTGCTACCTTGTGATAAAACTGTGCACACAA
TTATTCCATAGGAAATTTAATCAAAGTTTTTTAAAGTTAAAGTTAAAAACAAGTTGATAGCCAAAACTAGAATA
GTTAGCTAAACGAATTGAACACTCACAAAGGTGCCATAAAAAGCCTTAAATAGATTAACAAGAAACAAAATAA
CTAGTACAAATCCCTG

>Line C6
TCGCTGTTCGTCAGCCCGGCCTTGAATAATTCTTCTGTCATTGGCATTTTCACATTTCATTTAACTTTGGCCCGCT
CTGCTTTTGCTGCTGCTTCTGCTTTTTTGGGCCTTCGTAAGGCAGGACTAGGTTTTTGGTGCTTCTGTTGCCAAC
CGGAGCGGTAATCAGACTTCGGCCGAGGATTTTATCCGCTGCATTCCACAAATCGCTTGTTGGTTTTAGCGGC

CACAATCTGATAGTTTCACATTTGGTTAAATGTTCGTCATCAGCCAATCACTTTTGTCTGGTCCGCCAATGGGTT
TCAGCAGAAAGGGTTAGGTCATTTAATTTTATTGGAACTGAAATATACACAAACTTATGTAAAGAAAGATTTA
ACTTTTCTACTTGATTAAACTACTGATAATGTTTAAACTGAATTCTTTAAATACAAAATTCTATACAACAAATTCT
ATAGAAACGAATTTCGCTTACAATGCACATAGGTCTAGTAAATAACACTTTGACTACAAAGGCAAATTTCTAAA
AGAAAATGTACTAATTTCCTATACTTAAACTAAACCTGTACTTAAACTAGTGGCACTGAGATTCAAAACCAGGC
GCCAAAGCTTTCCCATCACTTATTGCAAGCAGTGCAAAGCTCCATGCAAGCAGTGCGACAAACTGTAGCCACA
TGGCCGGCCCCAAAAAAAAAGCAAAAAACAAAGCGAACAGCTGACTGAAACTTGCGATTTTCTGCTTCTGCAT
TTCGGCTTTGGCGCCCCGCGTTTGTTGTTTTGCTTATTATTCTGCGTTCTTCGCTCGCTGCGCGAATACCGAATT
CTGAACACTGAGACTGTGCCCGGCGCTCAAAAATCTTTCCCAAACTTTTCGGCATTTCCACGTGAATTTGAACG
GCCAAAGCACAAATCAAAAACATATTTTTTGAGAATTACCCGAGTGTCATAAATAGAGAGAGTGATAAGTCTT
TCTAAATGTTTAATAATTAGAGAATGCATAAACAAATGGTGGCATTTATTAGATGAAAATGCATAAAGACTTG
GCCAAAAATAAAACACGTAAAAAATCCTACATCATCTCTACTCTGGTTGCTACCTTGTGATAAAACTGTGCACA
CAATTATTCCATAGGAAATTTAATCAAAGTTTTTTAAAGTTAAAGTTAAAAACAAGTTGATAGCCAAAACTAGA
ATAGTTAGCTAAACGAATTGAACACTCACAAAGGTGCCATAAAAAGCCTTAAATA

**Supplementary Figure 1**

***Control experiment for adverse phenotypic effects of mifepristone***

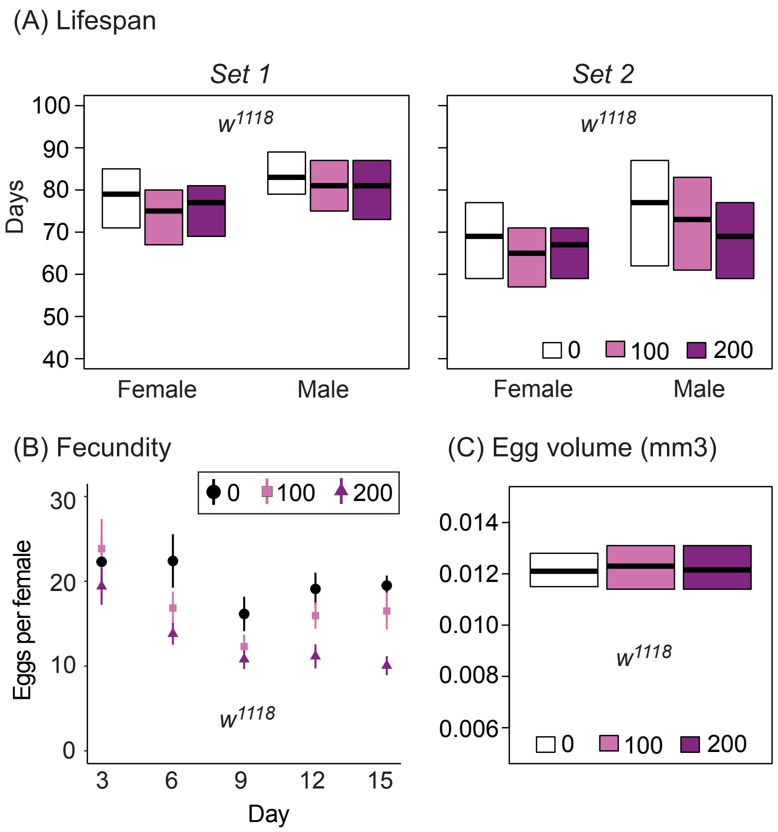


**Supplementary figure 1: No significant effects of mifepristone application on *da*-GS-GAL4/*w*$^{1118}$.**

Mifepristone application did not have a significant effect on lifespan (Batch 1 - female: $\chi^2$ = 6.5, *P* = 0.039, male: $\chi^2$ = 2.0, *P* = 0.37; Batch 2 - female: $\chi^2$ = 2.4, *P* = 0.3, male: $\chi^2$ = 10.5, *P* = 0.0052), nor on egg volume (*F* = 0.84, *P* = 0.44). A mifepristone concentration of 200 µg/ml did, however, have a significant effect on egg laying rates ($\chi^2$ = 28.9, *P* < 0.0001). The concentration of mifepristone was therefore limited to 100 µg/ml for the fecundity experiments ($\chi^2$ = 4.9, *P* = 0.085). The effects of mifepristone concentration (0, 100, or 200 µg/ml) were considered significant when P<0.0042 (Bonferroni: 0.05/12), whereas effects on egg laying rate and egg volume were considered significant when P<0.01 (Bonferroni: 0.05/5).

**Supplementary Table 1**

***Confirmation of knockdown of Eip75B by RNAi***

**Supplementary table 1:** The relative expression of *Eip75B* after mifepristone application (100 µg/ml or 200 µg/ml), was compared to the expression level in the control (no mifepristone) using quantitative real-time PCR.

| Acronym | Construct | *Eip75B* relative expression | | *F* | *P* |
|---|---|---|---|---|---|
| | | 100 µg/ml | 200 µg/ml | | |
| E01 | TRiP.GLC01418 | 0.86 | 0.82 | 6.11 | 0.020 |
| E03 | TRiP.JF02257 | 0.83 | 0.72 | 19.66 | 0.00013 |
| E10 | VDRC #108399 | 1.08 | 0.87 | 1.95 | 0.17 |
| E44 | VDCR #44851 | 0.85 | 0.82 | 4.51 | 0.043 |

**Supplementary Table 2**

***Estimation of genetic contribution of the Eip75B SNP to fecundity***

**Supplementary table 2:** The table below gives estimates from the F1 association study of the genotypic values of the three different genotypes (TT, TG, GG), the additive variances of fecundity attributable to the candidate SNP ($V_G$), and the intraclass correlation coefficient or "isofemale heritability" ($t$, i.e., the proportion of total variance attributable to the SNP). See the Methods section for details.

| | **day 3** | **day 6** | **day 9** | **day 12** | **day 15** | **day 18** | **day 21** | **day 24** | **day 27** | **day 30** |
|---|---|---|---|---|---|---|---|---|---|---|
| *Average effects of genotypes (= genotypic values, i.e., mean number eggs per female)* | | | | | | | | | | |
| **TT** | 25.3 | 26.8 | 23.1 | 26.8 | 21.2 | 19.3 | 19.0 | 17.5 | 19.3 | 15.7 |
| **TG** | 24.9 | 28.8 | 25.7 | 31.4 | 25.3 | 23.9 | 23.6 | 20.8 | 21.9 | 19.5 |
| **GG** | 24.2 | 27.7 | 26.3 | 33.1 | 27.4 | 23.4 | 23.5 | 21.7 | 23.4 | 21.3 |
| *$V_G$ and t calculated using allele frequencies* | | | | | | | | | | |
| $V_G$ | 0.12 | 0.91 | 1.91 | 6.29 | 5.46 | 5.13 | 5.20 | 3.15 | 2.17 | 4.56 |
| $t$ | 0.003 | 0.018 | 0.057 | 0.085 | 0.096 | 0.095 | 0.111 | 0.082 | 0.053 | 0.083 |
| *$V_A$ and t calculated using random effects model* | | | | | | | | | | |
| $V_G$ | 0.0 | 0.0 | 1.80 | 8.11 | 8.03 | 4.96 | 5.47 | 3.65 | 2.69 | 6.38 |
| $t$ | 0.000 | 0.000 | 0.054 | 0.107 | 0.135 | 0.092 | 0.116 | 0.093 | 0.064 | 0.112 |

**Supplementary Table 3**

***qRT-PCR primers used to quantify Eip75B knockdown***

**Supplementary table 3:** Overview of the primer sets used to quantify *Eip75B* and four reference genes with quantitative real-time PCR. The amplicon size (bp) and the primer efficiency of each set is given. Additional information on the qRT-PCR experiment in given in the Methods section.

| **Gene** | **Forward primer** | **Reverse primer** | **Amplicon (bp)** | **Primer efficiency*** |
|---|---|---|---|---|
| *Eip75B* | GGAGGATGTCCCCTGCGCAACC | GAGTATTCCGATCTGCTGCTGCTG | 168 | 1.88 |
| *TBP* | CGGTTTCCCTGCAAAGTTCCTCGA | CACGATTCGAGGTCGCACCATACG | 172 | 1.92 |
| *Cyp1* | GTCGGCAGCGGCATTTCAGAT | CTGCACGCTGACGAAGCTAGG | 78 | 1.91 |
| *Ef1a48D* | TGCCACACCGCTCACATTGCT | CACGCACAGGGGCTTAGAGG | 150 | 1.96 |
| *Rap2l* | TATTGGACACCGCGGGCACA | TGGGTGCTGGCTGACTTCCT | 168 | 1.96 |

**qPCR protocol: 95°C for 10 min; 40 cycles with 95°C for 15 sec and 60°C for 1 min; melting analysis from 60°C until 95°C (+0.05 °C/s).*

### Supplementary Table 4

#### *F1 crosses for SNP association study*

**Supplementary table 4:** Setup of F1 crosses for assessing the association between the focal SNP and lifespan and fecundity. The table indicates the line numbers of the DGRP lines from which females and males were taken to generate the F1 crosses with either “GG”, “TT”, or “TG” genotype.

| Genotype | Cross | Females | Males |
|---|---|---|---|
| GG | 1 | 142 | 787 |
| GG | 2 | 304 | 852 |
| GG | 3 | 319 | 379 |
| GG | 4 | 356 | 821 |
| GG | 5 | 386 | 324 |
| GG | 6 | 555 | 105 |
| GG | 7 | 73 | 377 |
| GG | 8 | 796 | 832 |
| GG | 9 | 83 | 790 |
| GG | 10 | 908 | 158 |
| TT | 11 | 239 | 362 |
| TT | 12 | 306 | 208 |
| TT | 13 | 340 | 375 |
| TT | 14 | 385 | 508 |
| TT | 15 | 391 | 853 |
| TT | 16 | 405 | 129 |
| TT | 17 | 426 | 566 |
| TT | 18 | 443 | 843 |
| TT | 19 | 716 | 355 |
| TT | 20 | 819 | 761 |

| Genotype | Cross | Females | Males |
|---|---|---|---|
| TG | 21 | 195 | 555 |
| TG | 22 | 158 | 42 |
| TG | 23 | 356 | 843 |
| TG | 24 | 712 | 83 |
| TG | 25 | 287 | 370 |
| TG | 26 | 812 | 589 |
| TG | 27 | 57 | 304 |
| TG | 28 | 900 | 852 |
| TG | 29 | 100 | 379 |
| TG | 30 | 822 | 492 |
| TG | 31 | 21 | 310 |
| TG | 32 | 461 | 517 |
| TG | 33 | 790 | 712 |
| TG | 34 | 324 | 391 |
| TG | 35 | 819 | 386 |
| TG | 36 | 375 | 217 |
| TG | 37 | 405 | 639 |
| TG | 38 | 821 | 721 |
